# SEQdata-BEACON: a comprehensive database of sequencing performance and statistical tools for performance evaluation and yield simulation in BGISEQ-500

**DOI:** 10.1101/652347

**Authors:** Yanqiu Zhou, Chen Liu, Rongfang Zhou, Anzhi Lu, Biao Huang, Liling Liu, Ling Chen, Bei Luo, Jin Huang, Zhijian Tian

**Affiliations:** BGI-Wuhan Clinical Laboratories, Wuhan, 430074, China

**Keywords:** BGISEQ-500, Prediction tools, Data analysis, Database

## Abstract

**Background:** BGISEQ-500 is based on DNBSEQ™ technology and superior in providing high outputs and requiring less cost. This sequencer has been widely used in various areas of scientific and clinical research. A better understanding of the sequencing process and sequencer performance is essential for stabilizing sequencing process, accurately interpreting sequencing results and efficiently solving sequencing troubles. To solve these problems, a comprehensive database SEQdata-BEACON was constructed to accumulate sequencing performance data in BGISEQ-500.

**Methods:** Totally 60 BGISEQ-500 sequencers in BGI-Wuhan lab were used to collect the sequencing performance data. Those lanes in paired-end 100 sequencing using 10bp barcode were chosen, and each lane containing 66 metrics was assigned a unique entry number as ID. The database was constructed in MySQL server 8.0 and the website was built on Apache (2.4.33 win64 VC15 server). The statistical analysis and linear regression models were generated by R program based on the data from November 2018 to April 2019.

**Results:** A total of 2236 entries were recorded in the database, including sample ID, yield, quality, machine state and supplies information. According to correlation matrix, the 52 numerical metrics were clustered into three groups signifying yield-quality, machine state and sequencing calibration. The metrics distributions also delivered some patterns and rendered clues for further explanation or analysis of the sequencing process. Using the data of total 200 cycles, the linear regression model well simulated the final outputs. Moreover, the predicted final yield could be provided in the 15^th^ cycle of the early stage of sequencing and the corresponding coefficient of determination R^2^ of the 200^th^ and 15^th^ cycle models were 0.97 and 0.81 respectively. The data source, statistical findings and application tools were all available in our website http://seqBEACON.genomics.cn:443/home.html. These resources can be used as a constantly updated reference for BGISEQ-500 users to comprehensively understand DNBSEQ™ technology, solve sequencing problems and optimize the sequencing process.

## Introduction

The next generation sequencing (NGS, also known as high-throughput sequencing) has led us into a genomic era. In the past 15 years, the development of sequencing technology was mainly committed to the reduction in cost and improvement in outputs, accuracy and read length. Nowadays, the sequencer manufacturers Illumina and Beijing Genomics Institute (BGI) provided high outputs and accuracy, while Pacific Bioscience and Oxford Nanopore offered long read length (Ansorge 2018; Goodwin et al. 2016). The first BGI sequencer BGISEQ-500 has launched in 2015 (http://en.mgitech.cn/), which was based on two key technologies: DNA nanoball (DNB) and Combinatorial Probe-Anchor Synthesis (cPAS). DNA library circles formed a single-stranded DNA molecule named as DNB by rolling circle amplification (RCA). The DNBs were dispersed and immobilized on a patterned array, then a probe was annealed to a DNA molecular anchor on the DNB. In each cycle, DNA polymerase incorporated one base labeled with fluorescence group, and the light signal was collected via a high-resolution imaging system and converted into bases after basecalling (Drmanac et al. 2010).

Recently, BGISEQ-500 platform has been widely used in a variety of sequencing types, such as whole genome sequencing (WGS), whole exome sequencing (WES), RNA-seq, small RNA and metagenomics (Chen et al. 2017; Fehlmann et al. 2016; Han et al. 2018; Huang et al. 2017; Xu et al. 2019). BGISEQ-500 sequencing platform has not only participated in transcriptome analysis of plant nitrogen metabolism and resistance response, but also in human clinical applications for the cancer genome sequencing and TP53 mutations detection in high-grade serous ovarian cancer (Chen et al. 2018; Li et al. 2018; Liu et al. 2018; Patch et al. 2018). In addition, it is reported that BGISEQ-500 had good performance in single cell resolution, such as scRNA-seq and scCAT-seq (Liu et al. 2019; Natarajan et al. 2019; Zhao et al. 2018). Compared with Illumina platforms, BGISEQ-500 demonstrated comparable SNP detection accuracy in WGS, similarly consistency in variation detection in WES, and high concordance in transcriptome and metagenomic studies (Fang et al. 2018; Patch et al. 2018; Xu et al. 2019; Zhu et al. 2018). Therefore, DNBSEQ™ technology had the advantages of cost-effective, high reproducibility and low duplication rate which provided a new choice for resolve issues in scientific research, agriculture, environment and clinical applications.

Among the sequencing performance metrics in BGISEQ-500, yield (eg., Reads, the number of DNBs recognized by the Basecall software) and quality (eg., Q30, the percentage of bases with an error rate below 0.001) were the most concerned. Besides, other metrics regarding chemical reaction and instrument state were also recorded in the sequencing summary. However, we still lack a profound understanding of the metrics meaning especial the connection between them. As one of the world’s largest sequencing service providers, BGI performed thousands of sequencing runs each year. Accompanied with enormous nucleotide data, massive sequencing performance data was highly valuable for illustrating this complicated process and troubleshooting. Unfortunately, this type of datasets or databases are rare currently and a comprehensive database is required to integrate the abundant sequencer performance data.

In this study, we designed a database SEQdata-BEACON to comprehensively collect sequencing performance data in BGISEQ-500 including the information of sample ID, yield, quality, machine state and supplies. We calculated Pearson’s correlation of 52 numerical metrics for hierarchical clustering and analyzed their distribution patterns. We also used linear regression to establish the yield simulation models to investigate the connection between yield correlated metrics, and attempted to predict the final yield in the early stage of sequencing. All the data and statistical analysis results were exhibited in our open access website. These resources can be an updating reference dataset for BGISEQ-500 users from enterprises or colleges to gain a deeper understanding of DNBSEQ™ technology.

## Materials and Methods

### Data Collection and Database Construction

We chose 60 BGISEQ-500 sequencers in BGI-Wuhan lab and collected all available files generated after chemical reaction and basecalling. We accumulated the lanes in PE100 sequencing since it is the major sequencing type in BGISEQ-500. Each lane was assigned a unique entry number as ID and other 65 metrics were evaluated and extracted to form a database ‘SEQdata-BEACON’. No detailed sample information was entered into the database to protect the privacy of customers. The database was constructed in MySQL server 8.0 and the architecture was shown in Fig. 1.

**Figure 1.**
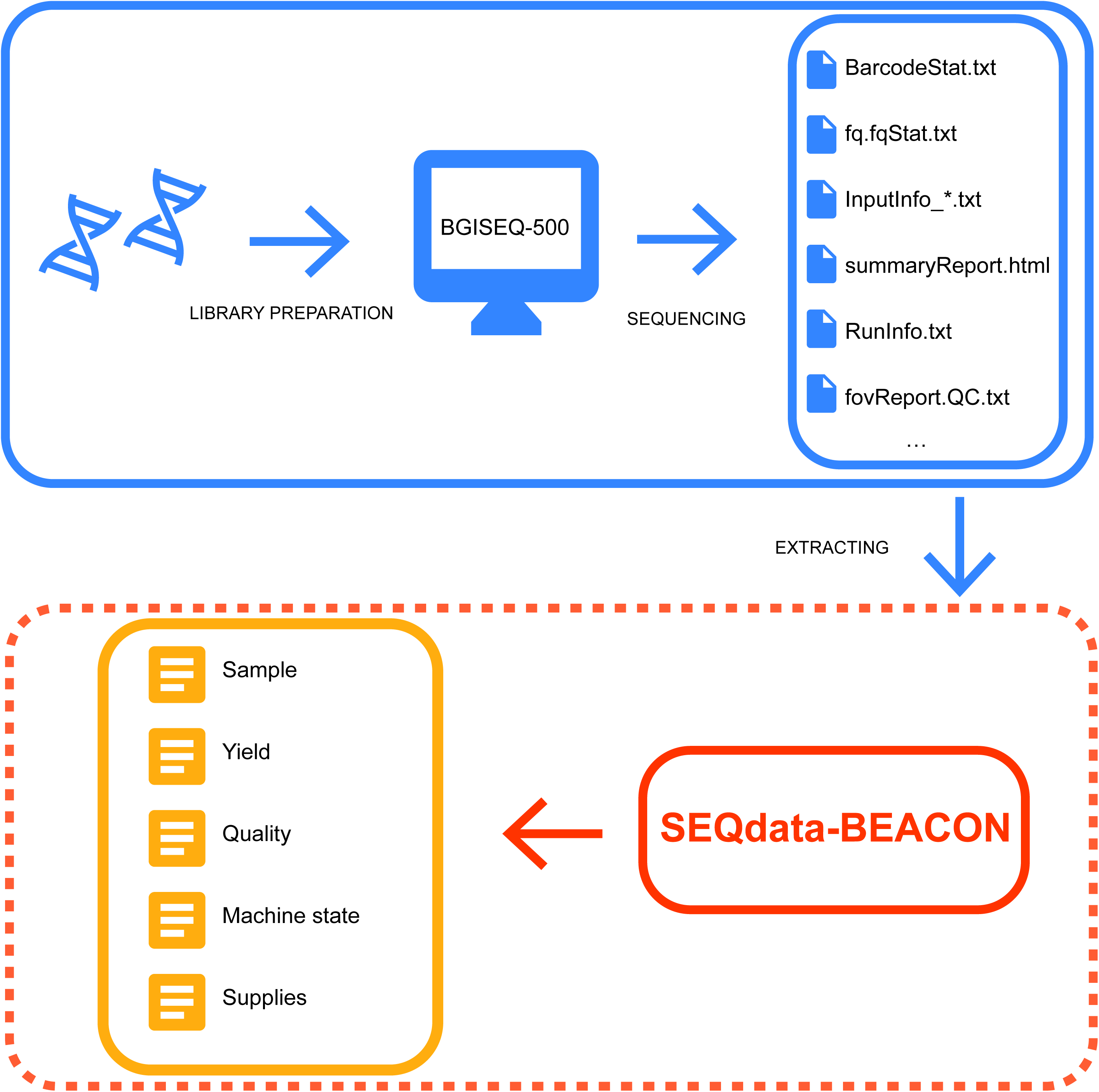
Schematic architecture of SEQdata-BEACON.

### Web Visual Interface and Statistical analysis

A user-friendly interface of SEQdata-BEACON was built on Apache (2.4.33 win64 VC15 server). An explorer Google Chrome 68.0.3440.106 was suggested to access the website. The statistical analysis and the figures in this study were generated based on the data from November 2018 to April 2019 by program R version 3.5.0 ×64 reserved the function to install modules/package.

### Yield simulation model

We tried to investigate the metrics correlated to yield and used them to simulate the final outputs. First, considering the sequencing principle, DNB was the decisive source of yield. The number of successfully fixed DNBs on the patterned array determined the ability to produce reads (Drmanac et al. 2010; Wang et al. 2019). TotalEsr (Total Effective Spot Rate) multiplied by Dnbnumber excluded those spots without DNB or with dark DNB not showing light signal, and represented the maximum signal that could be successfully collected. Second, some metrics that clustered with yield (Reads) may reflect the low-quality reads resulting from chemical reaction, light intensity, camera acquiring and basecalling in the sequencing process. At the end of sequencing, the sequencers filtered out these reads with low quality score which resulted in the loss of final yield. Thus, we constructed a linear regression model with TotalEsr*Dnbnumber, BIC, accGRR, SNR and FIT as variables of inputs, and output was yield (Reads). We used Turkey’s boxplot to pinpoint the possible outliers which defined as observations that fall below Q_1_-1.5 IQR or above Q_3_+1.5 IQR, and these outliers were excluded from the model construction. The LR model formula was as follows (Eqs. (1)):

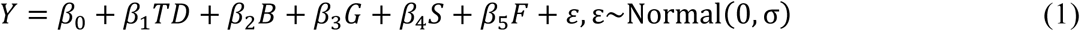

*Y* is the value of the dependent variable yield. *T* is the value corresponding to the current cycle in TotalEsr. *D* is the value of Dnbnumber at the beginning of the sequencing. *B, G, S* and *F* are the average values of BIC, accGRR, SNR and FIT for the first 200 cycles respectively. ε is observed error and obeys normal distribution. The model parameters *β*_*n*_ are the coefficient value and estimated using a regression model.

Further, we simulated the final yield based on the metric values in every 5 cycles. The formula was as follows (Eqs. (2)):

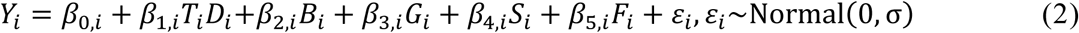

*Y*_*i*_ is the value of the dependent variable yield. *T*_*i*_ is the value corresponding to the current cycle i in TotalEsr. *D*_*i*_ is the value of Dnbnumber at the beginning of the sequencing. *B*_*i*_, *G*_*i*_, *S*_*i*_ and *F*_*i*_ are the average values of BIC, accGRR, SNR and FIT for the first i cycles respectively. *ε*_*i*_ is observed error in the i cycle and obeys normal distribution. The model parameters *β*_*n,i*_ are the coefficient value and estimated using a regression model. Both linear regression and backward elimination method of stepwise regression were conducted by program R version 3.5.0 ×64 reserved the function to install modules/package.

## Results

### Database Construction

DNA libraries were circularized, made into DNBs and loaded onto the patterned array. After successive chemical reaction, signal acquisition and basecalling, the sequencers normally generated a series of folders and files in each cycle or at the end of sequencing process. In BGISEQ-500, over 10 files recorded the information about the entire sequencing process, such as InputInfo_*.txt, RunInfo.txt, summaryReport.html, fovReport.QC.txt, BarcodeStat.txt and fq.fqStat.txt etc. The metrics values in these files were the preliminary dataset for ‘SEQdata-BEACON’, and totally 66 metrics were extracted and entered into our database (Fig. 1). From November 2018 to April 2019, ‘SEQdata-BEACON’ accumulated a total of 2236 entries in PE100 sequencing using 10bp barcode from 60 sequencers in BGI-Wuhan lab. The database primarily collected the information of sample ID, yield, quality, machine status, and supplies. The unique identifier for each entry, sequencing type and other sample information were stored as Sample ID. Regular quality control metrics such as Reads, Bases related to yield and Q30 related to quality were stored. Machine state such as Signal, Intensity and Theta were stored into the database. Sequencing reagents and sequencing time were also stored in the database.

### Web Visual Interface

To provide open access to our data, we designed a comprehensive website ‘SEQdata-BEACON’ with Home, Browse, Tools, Download and Guide pages to display the database and data-mining applications. The ‘Home’ page allowed the users to overview the introduction of our website and schematic architecture of our database (Fig. 2A). The ‘Browse’ page allowed the users to look through the numerical metric features including heatmap of Pearson’s correlation matrix and the distribution. For example, the distribution of FIT and its change per cycle were both manifested in charts for observing the distribution patterns and fluctuations (see the section ‘Statistical Finding: Metric Features’ for details). The ‘Tools’ page allowed the users to test our simulation model (Fig. 2B and C). Users could enter specified metric values in our website in the example format and click ‘Start’, then the expected yield confidential intervals will be shown (see the section ‘Statistical Finding: Yield simulation model’ for details). The ‘Download’ page allowed the users to obtain the data in EXCEL format according to our update time, while all the data and analysis results will be updated every two months. The ‘Guide’ page supplied a guideline for regular operation.

**Figure 2.**
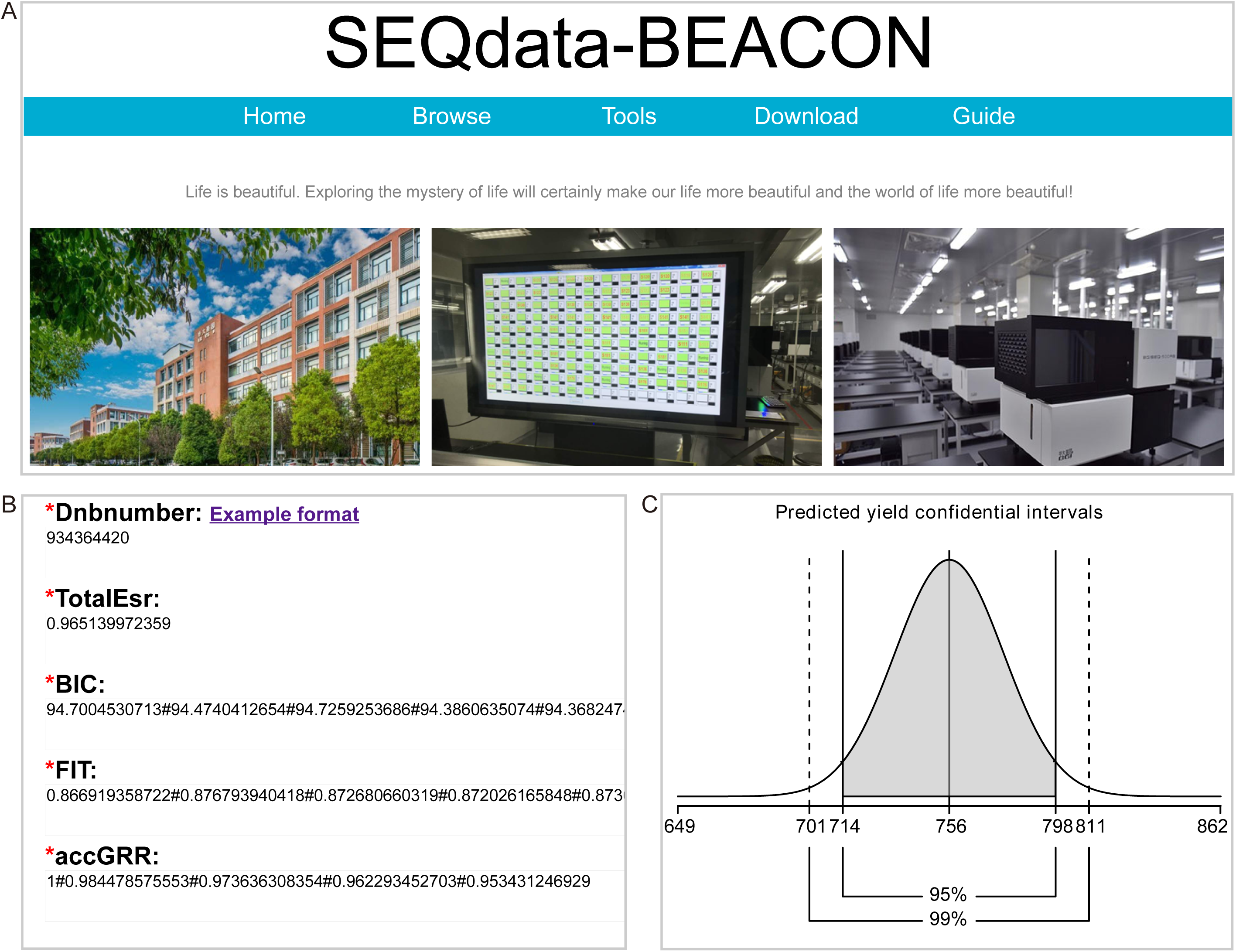
Web visual interface of SEQdata-BEACON. (A) “Home” page showed the database instruction and its five pages including Home, Browse, Tools, Download and Guide. (B) “Tools” page showed the input windows of metrics for users to test our yield simulation model. (C) “Tools” page showed and the resulting confidential intervals of predicted yield.

### Statistical Finding: Metric Features

Based on the data from November 2018 to April 2019, Pearson’s correlation of numerical metrics was calculated and the 52*52 correlation matrix was exhibited in the heatmap (Fig. 3). These 52 metrics were mainly clustered into three groups with 20, 15 and 17 metrics respectively, and the first group could be further divided into two branches which represented yield and quality. The other two groups represented machine state (e.g., Theta, tells the angle between the moving direction of the array stage and the track line on the array), and sequencing calibration (e.g., Lag, Runon, record the percentage of read strands being out of phase with the current cycle). Red blocks indicated for the positive correlation and blue blocks for the negative ones. It is shown that yield was positively correlated with quality, and the sequencing calibration were negatively correlated with them both. It is also denoted that some metrics were redundant, for example, Reads was highly correlated with other five yield related metrics, and metrics about CG contents were highly correlated with each other. Only one of the redundant metrics was reserved in further data analysis.

**Figure 3.**
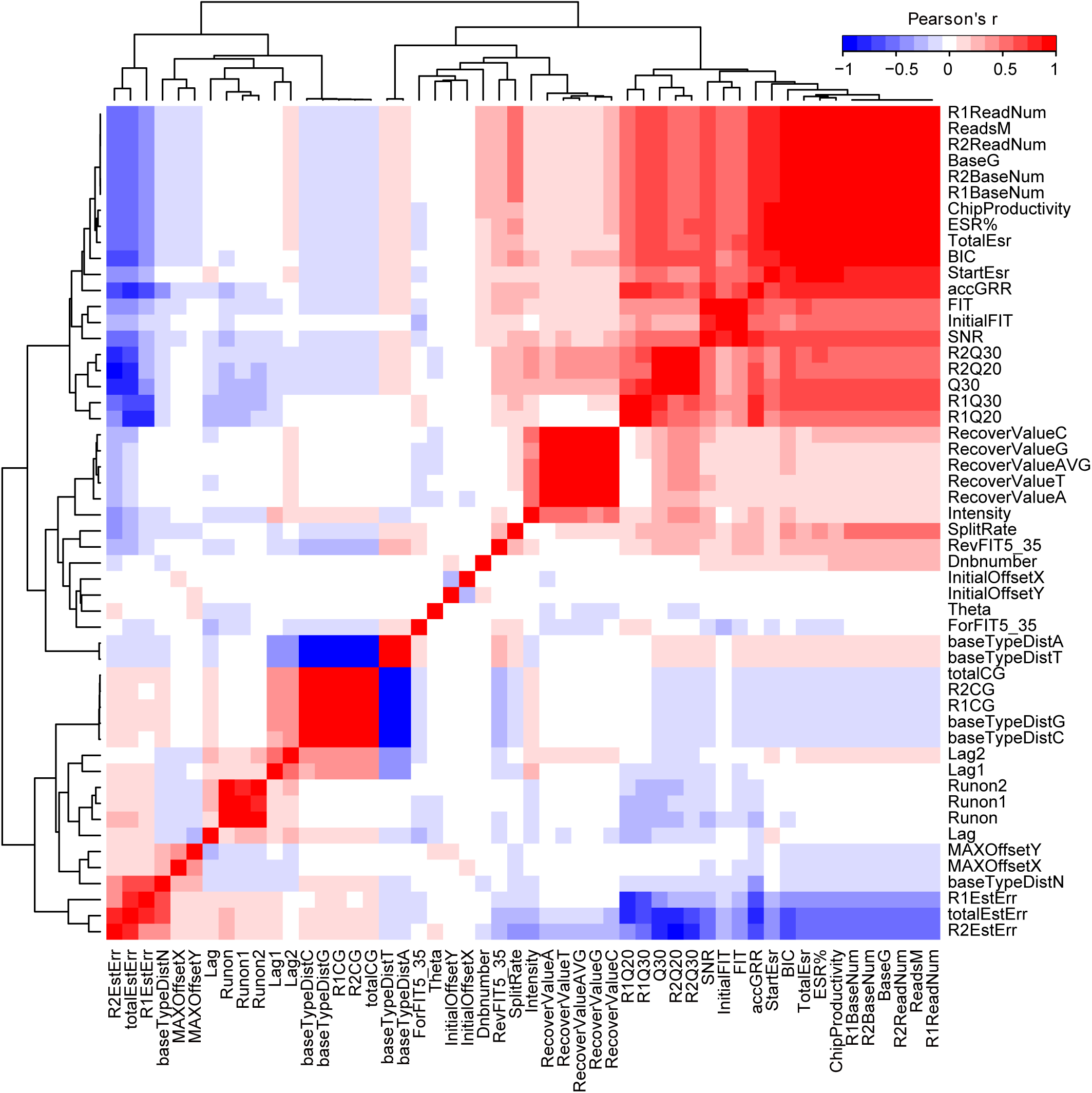
Numerical metrics correlation. Hierarchical clustering of the Pearson’s correlation matrix between 52 metrics.

Next, we investigated the distribution pattern of the above metrics. It is displayed in the scatterplot of Q30 versus Reads that yield was ranging from 350-850M, and Q30 was above 70%. The histogram on the upper and right part of chart also exhibited the distribution of Reads and Q30 respectively (Fig. 4A). According to the normal performance of BGISEQ-500, Reads was greater than 650M and Q30 was over 85% in each lane (http://en.mgitech.cn/). In the scatter plot, totally 2026 lanes located in the normal range, and the proportion of outliers was less than 10%, which suggested that the instrument performance was relatively stable. Furthermore, the distribution pattern of FIT was manifested in a histogram, and it was mostly around 0.80 (Fig. 4B). FIT value was presented cycle by cycle, and it was found to slowly decrease from 0.811 to 0.757 in read1 and from 0.835 to 0.763 in read2 with more deviation in the beginning to less deviation in the end of each read (Fig. 4C). The metric FIT suggested the distribution of differences between signal and noise for each base and the performance of optical path during the sequencing process (Wang et al. 2019). The deviation in each cycle may reflect the changing pattern of FIT in the sequencing process and hint the signal or optical path status indirectly.

**Figure 4.**
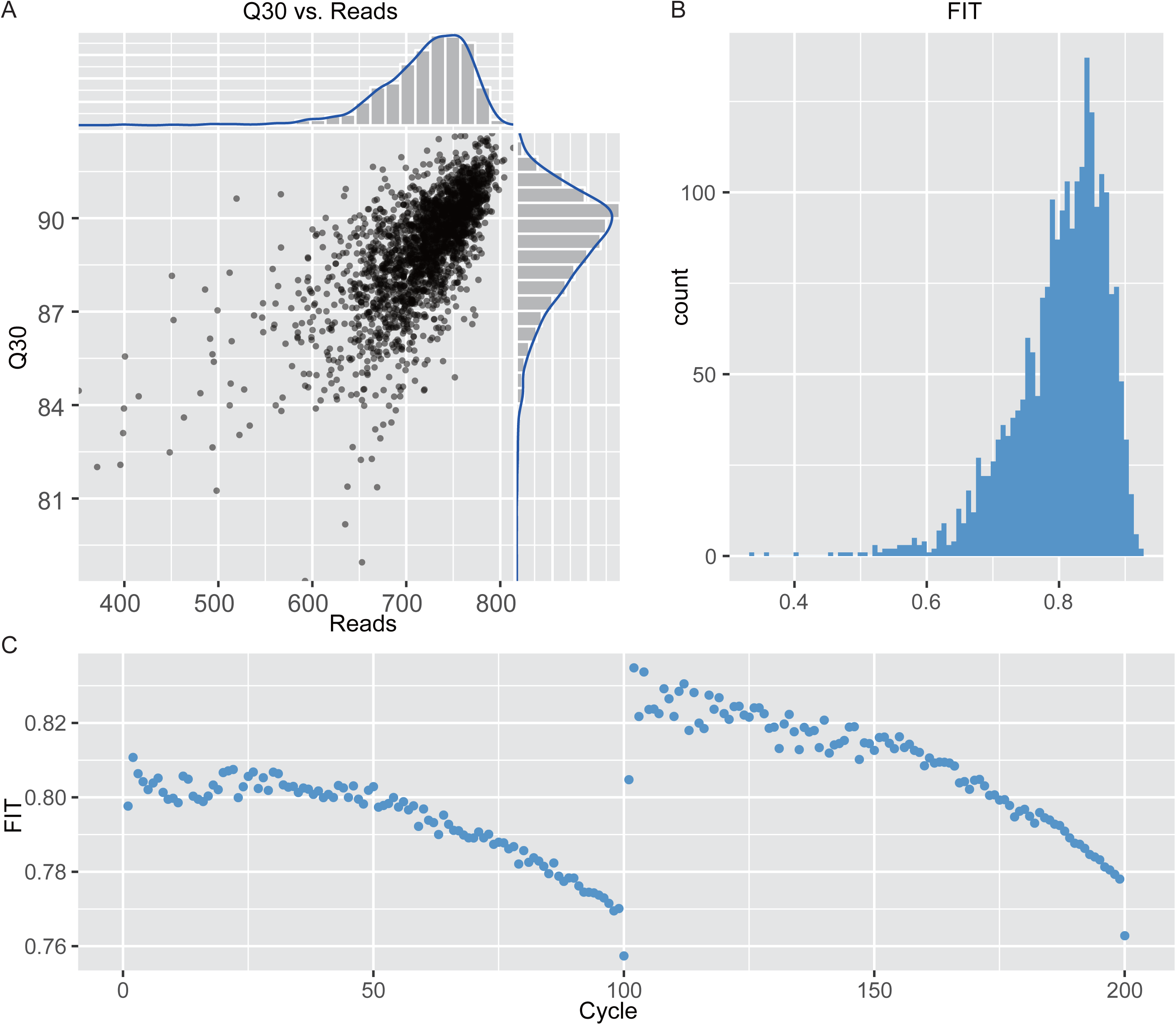
Distribution of Metrics in SEQdata-BEACON. (A) Q30 versus Reads, histogram of Q30 and Reads shown in grey, density profiles shown in blue. (B) Histogram of FIT in all entries. (C) Scatter plot of FIT through 200 cycles.

### Statistical Finding: Yield simulation model

In this study, considering the BGISEQ-500 sequencing principle and Pearson’s correlation matrix clustering results, the linear regression model included the variables TotalEsr, Dnbnumber, BIC, accGRR, SNR and FIT as predictors. Statistical program R was applied to construct the model using the data from November 2018 to April 2019. The LR model was Eq. (3) and Table 1 displayed the standard error, t-value and p-value of this function in regression results. It is indicated that the contribution of accGRR was not significant. Subsequently, backward elimination method of stepwise regression was used to build the model to find an optimum solution and the final LR model was Eq. (4). The regression results of this function pointed out that the contributions of TotalEsr*Dnbnumber, BIC, SNR, FIT were significant at the 1% probability level (Table 1). Here we compared the yield with predicted results, taking the coefficient of determination (R^2^) range from 0 to 1 to measure the accuracy of our model, which showed the value for R^2^ was 0.97 (Fig. 5). It is suggested that five metrics TotalEsr*Dnbnumber, BIC, SNR, FIT successfully simulated yield in a LR model.

**Table 1.** Model summaries of regressions linear for prediction of yield outputs.

| Independent variables | Outputs in Eq. (3) |  |  | Outputs in Eq. (4) |  |  | Outputs in Eq. (5) |  |  |
| --- | --- | --- | --- | --- | --- | --- | --- | --- | --- |
|  | Std. error | t-Value | P-value | Std. error | t-Value | P-value | Std. error | t-Value | P-value |
| Constant | 11.424 | -11.353 | < 2e-16** | 10.662 | -12.173 | < 2e-16** | 39.411 | -3.578 | 0.00035** |
| TotalEsr*Dnbnumber | 0.010 | 95.192 | < 2e-16** | 0.010 | 96.130 | < 2e-16** | 0.033 | 38.837 | < 2e-16** |
| BIC | 0.192 | 7.278 | 4.73e-13** | 0.178 | 7.873 | 5.46e-15** | 0.724 | 6.556 | 6.91e-11** |
| accGRR | 6.175 | 0.023 | 0.981 | --- | --- | --- | 59.009 | -10.243 | < 2e-16** |
| SNR | 0.388 | -8.586 | < 2e-16** | 0.388 | -8.596 | < 2e-16** | 0.653 | -5.307 | 1.23e-07** |
| FIT | 4.757 | 5.675 | 1.57e-08** | 4.577 | 5.905 | 4.08e-09** | 11.249 | 2.567 | 0.010* |

\*\*Significant at 1% probability level.
\* Significant at 5% probability level.
TotalEsr: ESR (Effective Spot Rate), the percentage of filtered Reads among the DNBs recognized by Basecalling. $ESR = \text{Total Reads} / \text{theoretical maximum reads number of one sequencing lane}$ . TotalEsr calculated ESR value in the first 15 cycles in read1 and read2, and kept constant in the rest of each read.
Dnbnumber: The theoretical maximum number of DNBs on the patterned array.
BIC: Basecall information content, the percentage of DNBs that can be used for Basecalling among the DNBs recognized by the optical system. $BIC = (\text{numbers of DNB that can be used for Basecalling} / \text{numbers of DNB that can be recognized by the optical systems}) \times 100\%$ .
accGRR: Accumulated Good Reads Rate, taking chastity greater than 0.6 as the filtering criteria, the percentage of filtered Reads among the DNBs recognized by Basecalling. $accGRR = \text{Total Reads} / \text{theoretical maximum reads number of one sequencing lane}$ . This value is only a statistical indicator which reflects the overall quality of the read (multi-cycle state).
SNR: Signal to Noise Ratio, taking the SNR calculation of a single DNB as an example, base A (maximum light intensity) is used as the signal, the CGT is the background, and the variance of the CGT light intensity is noise. $A\_SNR = A\_mean / CGT\_dev$ .
FIT: FIT value represents the distribution of differences between signal and noise for each base. The FIT value is higher when the
distribution of differences between signal to noise for each channel/color are more concentrated.

**Figure 5.**
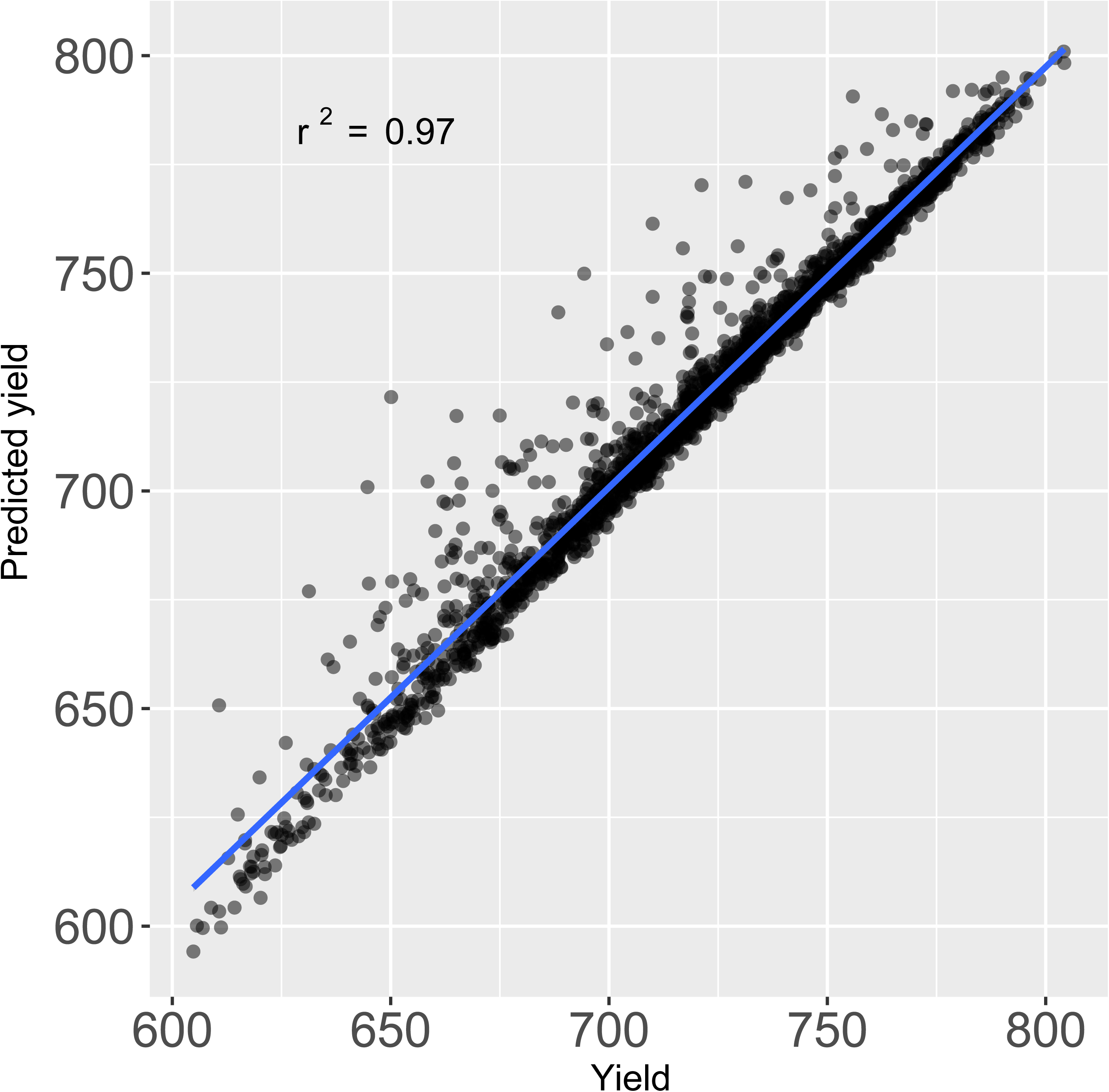
Cross-correlation of yield between predicted and actual values. A linear regression line is shown in blue line.

In addition, we hoped that this simulation model can be used to predict the final yield at the early stage of sequencing. In the sequencing process, the yield was assumed to be a function of TotalEsr, Dnbnumber, BIC, accGRR, SNR and FIT at the current running cycle. The model was conducted every 5 cycles using backward elimination regression by statistical program R, and totally 40 models were established from the cycle 1 to 200. Comparing the residual deviation of these models, the residuals of prediction models were fluctuated in the beginning of read1 and read2, which was mainly due to the establishment of the algorithm matrix by the sequencer (Fig. S1). However, the value for R^2^ of model in the 15^th^ cycle was 0.81, indicating that the final yield could be effectively simulated at this stage. The corresponding LR model was Eq. (5).

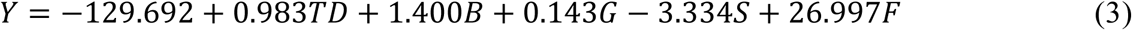

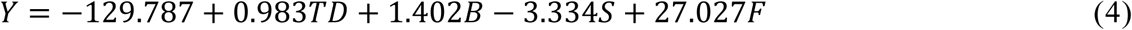

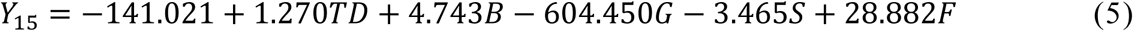

## Discussion

Continuous improvement in sequencing technologies and sequencers has provided us the choice to producing higher throughput data with less sequencing cost. With the upgrade of DNBSEQ™ technology, BGI has introduced more efficient sequencing platforms, such as MGISEQ-2000 and MGISEQ-T7. Meanwhile, because of the excellent performance of DNBSEQ™ technology, this series of sequencers had been widely used both in science research and clinical aspects. The sequencer performance metrics brought us a wealth of information of the sequencer, but our understanding of this information was seriously lacking. To gain insight into the sequencing performance in BGISEQ-500, we constructed a database ‘SEQdata-BEACON’ which has already contained 2236 entries with 66 metrics and comprehensively recorded the information of sample ID, yield, quality, machine state and supplies. The method of automatically collecting metric values from sequencing summary could effectively lighten human labor work, shorten time cost and improve data accuracy.

At present, in our 60 BGISEQ-500 sequencers in BGI-Wuhan lab, sequencing libraries were derived from plant, animal, microbial and human DNA or RNA samples, while PE100 sequencing using 10 bp barcode was for WGS, WES and RNA-seq. Therefore, the data we collected including libraries from most species and major types of sequencing applications. By analyzing the distribution characteristics of the numerical metrics, in Q30 versus Reads scatterplot, 90.6% lanes had reads more than 650M and Q30 above 85% which proved that BGISEQ-500 was stable and reliable in the massively parallel sequencing. Without the risk of index hopping, DNBSEQ™ could provide excellent sequencing data with less duplication and error, which could have extensive application in population scale sequencing projects, such as the 10KP (10,000 Plants) Genome Sequencing Project (Cheng et al. 2018; Li et al. 2019).

In order to study the correlation of yield-correlated metrics, we used the backward elimination method of stepwise regression to establish a yield simulation model with the value for R^2^ was 0.97. The model presented good simulation, which suggested that TotalEsr, Dnbnumber, BIC, SNR and FIT were contributed to the yield. While predicting the final production, we still used all six parameters to construct 40 prediction models using stepwise regression. From the residual deviation of the models, it is found that the final yield can be predicted in the 15^th^ cycle at the early stage of sequencing, and the small change in the residual within read1 and read2 implied the little fluctuation of the metrics during the sequencing process. The study of yield related metrics and their relationship began with constructing linear regression model which is a common statistical technique for simulating the associations between variables. However, it is not excluded that neural network models, machine learning model or other methods which may be utilized to get better simulation results. Furthermore, we wanted to investigate the quality associated metrics and establish a quality simulation model. Combined with yield simulation model, these two models may effectively simulate the sequencing results and bring us more ideas to increase the sequencing performance. With the updating of the database and website every two months, it is expected that the accumulation of data, enrichment of metrics together with the upgrade of algorithms will develop more data-mining applications.

We have established the first reported BGISEQ-500 sequencing performance database and website to comprehensively collect and summarize performance data. We paid more attention to the data accumulation, and hope to explore the data patterns by statistical analysis and interpret sequencing results. The database presented data and statistical results through the website, providing an ongoing reference dataset which gave users opportunities to promptly understand the performance and advantages of the sequencer. Recently, some clues were found in monitoring the instrument degradation by calculating error rates in Nextseq 500 (Manley et al. 2016). We also expected that our charts of metrics distributions could imply the troubles in the sequencing process. Recent investigations reported that MGISEQ-2000 had comparable SNP detection accuracy in WGS and high gene detection in scRNA-seq compared with Illumina platforms (Gorbachev et al. 2019; Senabouth et al. 2019). In the future, we plan to gather more sequencing platforms of DNBSEQ™ technology, which will provide an integrated statistical reference for BGI sequencers, and is beneficial to fully understand the sequencer performance of this series. Moreover, we also hope to add sequencer PacBio Sequel II and Oxford Nanopore PromethION to get deeper understanding of single-molecule sequencing technology. We expected SEQdata-BEACON to be a comprehensive platform: with data accumulation and data-mining, it could present the performance of the sequencing platform from various aspects; by adding functional modules such as QC metrics models and metrics standards, it also could provide users useful suggestions for optimization and troubleshooting to solve their problems.

## Conclusion

Widespread application of NGS resulted in large amount of data, including nucleotide sequences and sequencing process performance. We designed a database SEQdata-BEACON to accumulate sequencing performance data in BGISEQ-500 containing 66 metrics of sample ID, yield, quality, machine state and supplies. The correlation matrix of 52 numerical metrics was clustered into three groups covering the information of yield-quality, machine state and sequencing calibration. The distribution of numerical metrics presented some features and provided clues for further interpreting the metric meanings and analysis. We also constructed linear regression models to well simulate the final yield using metrics values in 200^th^ and 15^th^ cycle. The data source, statistical findings and application tools were all available in our website (http://seqBEACON.genomics.cn:443/home.html), which can facilitate BGISEQ-500 users from enterprises or colleges to understand and interpret their sequencing results.

## Acknowledgments

We would like to acknowledge the ongoing contributions and support of all our BGI employees.

We thank Zetao Bai (Oil Crops Research Institute, Chinese Academy of Agriculture Sciences, Wuhan, China) for her assistance editing this manuscript.

## Additional Information and Declarations

### Author Contributions

Yanqiu Zhou, Chen Liu and Rongfang Zhou constructed the database, analyzed the data, prepared figures and/or tables, drafted the work or revised it critically for important content. Anzhi Lu Biao Huang, Liling Liu, Ling Chen, Bei Luo contributed reagents/materials/analysis tools.

Jin Huang, Zhijian Tian conceived and designed the experiments, approved the final draft of the manuscript submitted for review and publication.

## Supplemental Information

**S1 Fig. Residual deviation of LR model every 5 cycles.** Box plot displayed the residuals of all the 40 LR models, each box manifested the median and first and third quartiles, star means the standard deviation.

**Figure S1. Residual deviation of LR model every 5 cycles.** Box plot displayed the residuals of all the 40 LR models, each box manifested the median and first and third quartiles, star means the standard deviation.

