## Supplementary figures and images for "SEQdata-BEACON: a comprehensive database of sequencing performance and statistical tools for performance evaluation and yield simulation in BGISEQ-500"

### Supplemental Figure S1

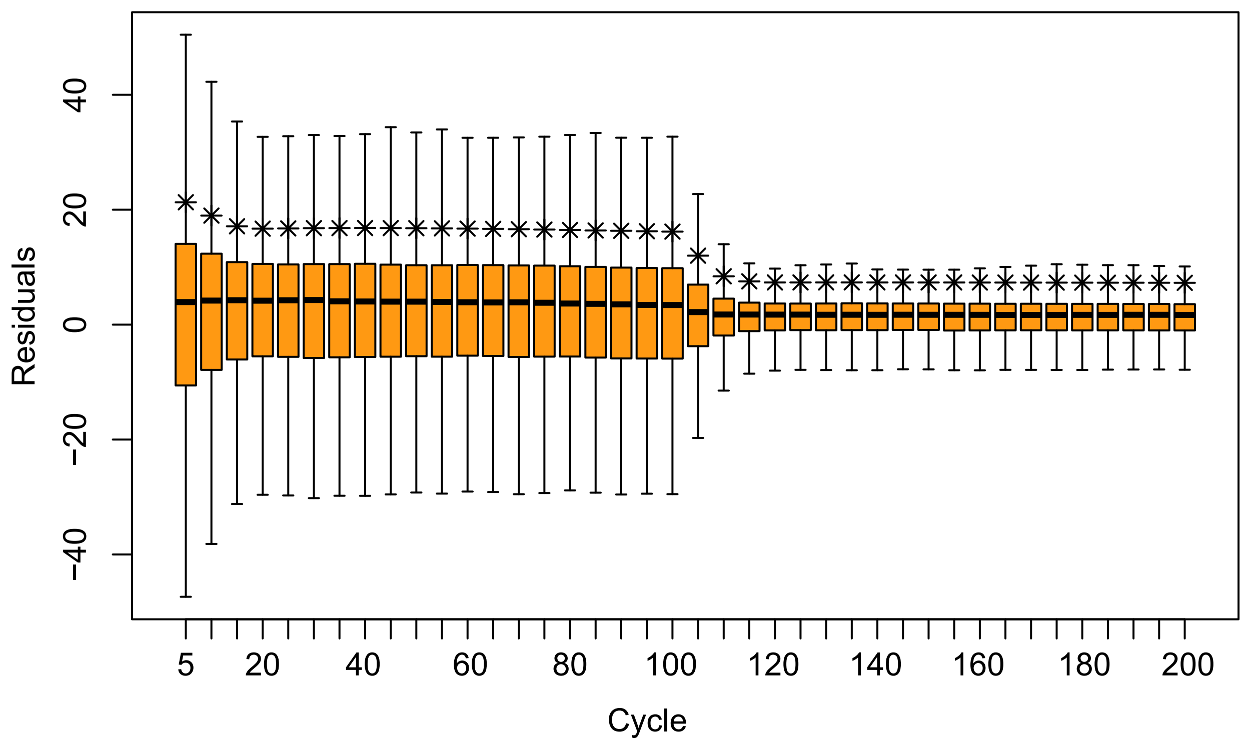
